# Neural Field Models for Latent State Inference: Application to Large-Scale Neuronal Recordings

**DOI:** 10.1101/543769

**Authors:** M. E. Rule, D. Schnoerr, M. H. Hennig, G. Sanguinetti

## Abstract

Large-scale neural recording methods now allow us to observe large populations of identified single neurons simultaneously, opening a window into neural population dynamics in living organisms. However, distilling such large-scale recordings to build theories of emergent collective dynamics remains a fundamental statistical challenge. The neural field models of Wilson, Cowan, and colleagues remain the mainstay of mathematical population modeling owing to their interpretable, mechanistic parameters and amenability to mathematical analysis. Inspired by recent advances in biochemical modelling, we develop a method based on moment closure to interpret neural field models as latent state-space point-process models, making them amenable to statistical inference. With this approach we can infer the intrinsic states of neurons, such as active and refractory, solely from spiking activity in large populations. After validating this approach with synthetic data, we apply it to high-density recordings of spiking activity in the developing mouse retina. This confirms the essential role of a long lasting refractory state in shaping spatio-temporal properties of neonatal retinal waves. This conceptual and methodological advance opens up new theoretical connections between mathematical theory and point-process state-space models in neural data analysis.

**Significance:** Developing statistical tools to connect single-neuron activity to emergent collective dynamics is vital for building interpretable models of neural activity. Neural field models relate single-neuron activity to emergent collective dynamics in neural populations, but integrating them with data remains challenging. Recently, latent state-space models have emerged as a powerful tool for constructing phenomenological models of neural population activity. The advent of high-density multi-electrode array recordings now enables us to examine large-scale collective neural activity. We show that classical neural field approaches can yield latent state-space equations and demonstrate that this enables inference of the intrinsic states of neurons from recorded spike trains in large populations.

## 1 Introduction

Neurons communicate using electrical impulses, or spikes. Understanding the dynamics and physiology of collective spiking in large networks of neurons is a central challenge in modern neuroscience, with immense translational and clinical potential. Modern technologies such as high-density multi-electrode arrays (HDMEA) enable the simultaneous recording of the electrical activity of thousands of interconnected neurons, promising invaluable insights into neural dynamics at the network level. However, the resulting data is high-dimensional and frequently exhibits complex, non-linear dynamics, presenting formidable statistical challenges.

Due to the high complexity of the data, most analyses of neuronal population activity take a descriptive approach, adopting methods from statistical signal processing such as state-space models (SSM; Paninski et al. 2010; Zhao and Park 2016, 2017; Sussillo et al. 2016; Aghagolzadeh and Truccolo 2016; Linderman et al. 2016; Gao et al. 2016) or autoregressive generalized-linear point-process models (PP-GLM; Paninski 2004; Pillow et al. 2008; Truccolo et al. 2005; Truccolo 2016). Such methods capture the population statistics of the system, but fail to provide mechanistic explanations of the underlying neural dynamics. While this phenomenological description is valuable and can aid many investigations, the inability to relate microscopic single-neuron properties to emergent collective dynamics limits the scope of these models to extract biological insights from these large population recordings.

Connecting single-neuron dynamics with population behavior has been the central focus of research within the theoretical neuroscience community over the last four decades. Neural field models (Amari, 1977; Wilson et al., 1972; Cowan, 2014; Bressloff, 2012) have been crucial in understanding how macroscopic firing dynamics in populations of neurons emerge from the microscopic state of individual neurons. Such models have found diverse applications including working memory (see Durstewitz et al. 2000 for a review), epilepsy (e.g. Zhang and Xiao 2018; Proix et al. 2018; González-Ramírez et al. 2015; Martinet et al. 2017), and hallucinations (e.g. Ermentrout and Cowan 1979; Bressloff et al. 2001; Rule et al. 2011), and have been successfully related to neuroimaging data such as Electroencepelography (EEG; Moran et al. 2013; Bojak et al. 2010; Pinotsis et al. 2012), Magnetoencephelography (MEG; Moran et al. 2013), electromyography (EMG; Nazarpour et al. 2012), and Functional Magnetic Resonance Imaging (fMRI; Bojak et al. 2010), which measure average signals from millions of neurons. Nevertheless, using neuralfield models to model HDMEA spiking data directly remains an open statistical problem: HDMEA recordings provide sufficient detail to allow modeling of individual neurons, yet the large number of neurons present prevents the adoption of standard approaches to non-linear data assimilation such as likelihood free inference.

In this paper, we bridge the data-model divide by developing a statistical framework for Bayesian modeling in neural field models. We build on recent advances in stochastic spatio-temporal modeling, in particular a recent result by Schnoerr et al. (2016) which showed that a spatio-temporal agent-based model of reaction-diffusion type, similar to the ones underpinning many neural field models, can be approximated as a spatio-temporal point process associated with an intensity (i.e. density) field that evolves in time. Subsequently, Rule and Sanguinetti (2018) illustrated a moment-closure approach for mapping stochastic models of neuronal spiking onto latent state-space models, preserving the essential coarse-timescale dynamics. Here, we demonstrate that a similar approach can yield state-space models for neural fields derived directly from a mechanistic microscopic description. This enables us to leverage large-scale spatio-temporal inference techniques (Cseke et al., 2016; Zammit-Mangion et al., 2012) to efficiently estimate an approximate likelihood, providing a measure of fit of the model to the data that can be exploited for data assimilation. Our approach is in spirit similar to latent variable models such as the Poisson Linear Dynamical System (PLDS; Macke et al. 2011; Aghagolzadeh and Truccolo 2016; Smith and Brown 2003), with the important difference that the latent variables reflects non-linear neural field dynamics that emerge directly from a stochastic description of single-neuron activity (Bressloff, 2009; Buice et al., 2010; Touboul and Ermentrout, 2011).

We apply this approach to HDMEA recordings of spontaneous activity from ganglion cells in the developing mouse retina (Maccione et al., 2014), showing that the calibrated model effectively captures the non-linear excitable phenomenon of coordinated, wave-like patterns of spiking (Meister et al., 1991) that have been considered in both discrete (Hennig et al., 2009a) and continuous neural-field models before (Lansdell et al., 2014).

## 2 Results

### 2.1 High level description of the approach

We would like to explain large-scale spatio-temporal spiking activity in terms of the intrinsic states of the participating neurons, which we cannot observe directly. Latent state-space models (SSMs) solve this problem by describing how the unobserved states of neurons relate to spiking observations, and predict how these latent states evolve in time. In this framework, one estimates a distribution over latent states from observations, and uses a forward model to predict how this distribution evolves in time, refining the latent-state estimate with new observations as they become available. This process is often called ‘data assimilation’. However, in order to achieve statistical tractability, SSMs posit simple (typically linear) latent dynamics, which cannot be easily related to underlying neuronal mechanisms. Emergent large-scale spatio-temporal phenomena such as travelling waves typically involve multiple, coupled populations of neurons and nonlinear excitatory dynamics, both of which are difficult to incorporate into conventional state-space models.

Fortunately, mathematical neuroscience has developed methods for describing such dynamics using neural field models. Neural field models map microscopic dynamics to coarse-grained descriptions of how population firing rates evolve. This provides an alternative route to constructing latent state-space models for large-scale spatio-temporal spiking datasets. However, neural field models traditionally do not model statistical uncertainty in the population states they describe, which makes it difficult to deploy them as statistical tools to infer the unobserved, latent states of the neuronal populations. A model of statistical uncertainty is important for describing the uncertainty in the estimated latent states (posterior variance), as well as correlations between states or spatial regions. As we will illustrate, work over the past decades to address noise and correlations in neural field models also provides the tools to employ such models as latent SSMs in data-driven inference.

At a high level then, our approach follows the usual derivation of neural field models, starting with an abstract description of single-neuron dynamics, and considers how population averages evolve in time. Rather than deriving a neural-field equation for the population mean rate, we instead derive two coupled equations for the mean and covariance of population states. We interpret these two moments as a Gaussian-process estimate of the latent spatio-temporal activity, and derive updates for how this distribution evolves in time and how it predicts spiking observations. This provides an interpretation of neural-field dynamics amenable to state-space inference, which allows us to infer neural population states from spiking observations.

### 2.2 Neural field models for refractoriness-mediated retinal waves

Most classical neural-field models (Wilson and Cowan, 1972, 1973) consider two neuron states: neurons may be either actively spiking (*A* state), or quiescent (*Q* state). However, voltage and calcium gated conductances typically lead to refractory states, which can be short following individual spikes, or longer after more intensive periods of activity. An excellent example of the importance of a refractory mechanism is found in the developing retina, where a slow afterhyperpolarization (sAHP) current mediates the long-timescale refractory effects that strongly shapes the spatio-temporal dynamics of spontaneous retinal waves (Hennig et al., 2009b). To address this, we explicitly incorporate additional refractory (*R*) states into our neural field model (e.g. Buice and Cowan 2007, 2009; Figure 1). In the following, we first outline a non-spatial model for such system, before extending it to a spatial setting with spatial couplings. Finally, we develop a Bayesian inference scheme for inferring latent states from observational data.

**Figure 1:**
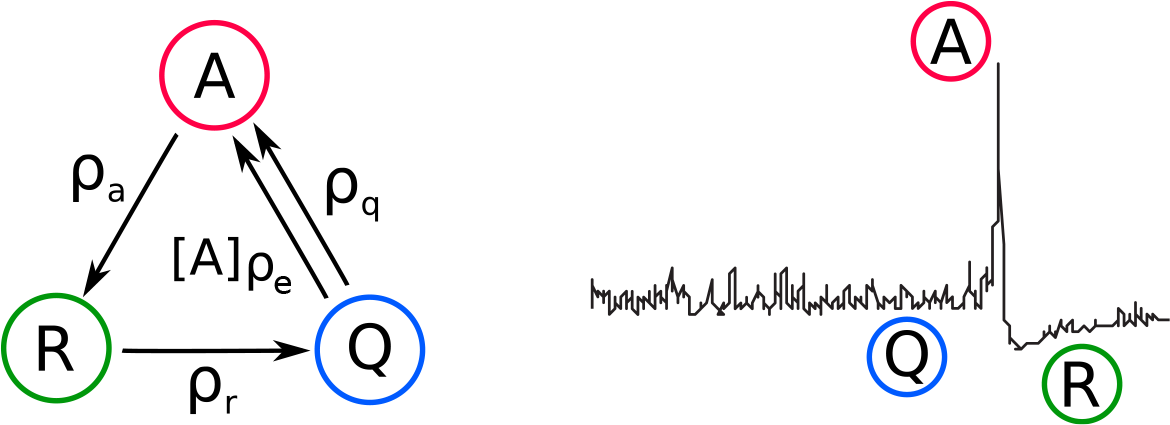
3-state Quiescent-Active-Refractory (QAR) neural-field model. Cells in the developing retina are modeled as having three activity states. Active cells (*A*; red) fire bursts of action potentials, before becoming refractory (*R*; green) for an extended period of time. Quiescent (*Q*; blue) cells may burst spontaneously, or may be recruited into a wave by other active cells. These three states are proposed to underlie critical multi-scale wave dynamics (Hennig et al., 2009b).

### 2.3 A stochastic three-state neural mass model

We now consider the neural field model with three states as a generic model of a spiking neuron (Figure 1), where a neuron can be in either an actively spiking (A), refractory (R), or quiescent (Q) state. We assume that the neurons can undergo the following four transitions:

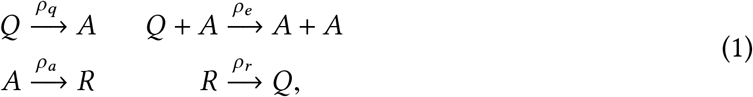

i.e. quiescent neurons transition spontaneously to the active state; active neurons excite quiescent neurons; active neurons become refractory, and refractory neurons become quiescent. The *ρ*_(·)_ denote corresponding rate constants.

For illustration, we first consider the dynamics of a local (as opposed to spatially-extended) population of neurons. In this case the state of the system is given by the non-negative number counts *Q*, *A* and *R* of the respective neuron types (we slightly abuse notation here and use *Q*, *A*, and *R* both as symbols for the neuron states and as variables counting the neurons in the corresponding states; see Figure 2 for an illustration). The time evolution of the corresponding probability distribution to be in a state (*Q*, *A*, *R*) at a certain time point is then given by a master equation (Buice and Cowan 2007; Ohira and Cowan 1993; Bressloff 2009; Methods: *Moment-closure for a single population*). Due to the nonlinear excitatory interaction *Q*+*A*→*A*+*A* in Eq. (1), no analytic solutions to the master equation are known. To get an approximate description of the dynamics, we employ the Gaussian moment closure method which approximates the discrete neural counts (*Q*, *A*, *R*) by continuous variables, and assumes a multivariate normal distribution (Figure 2B; Goodman 1953; Whittle 1957; Gomez-Uribe and Verghese 2007; Bressloff 2009; Buice et al. 2010; Schnoerr et al. 2017; Rule and Sanguinetti 2018). This allows one to derive a closed set of ordinary differential equations for the mean and covariance of the approximate process which can be solved efficiently numerically (Methods: *Moment-closure for a single population*; Figure 2).

**Figure 2:**
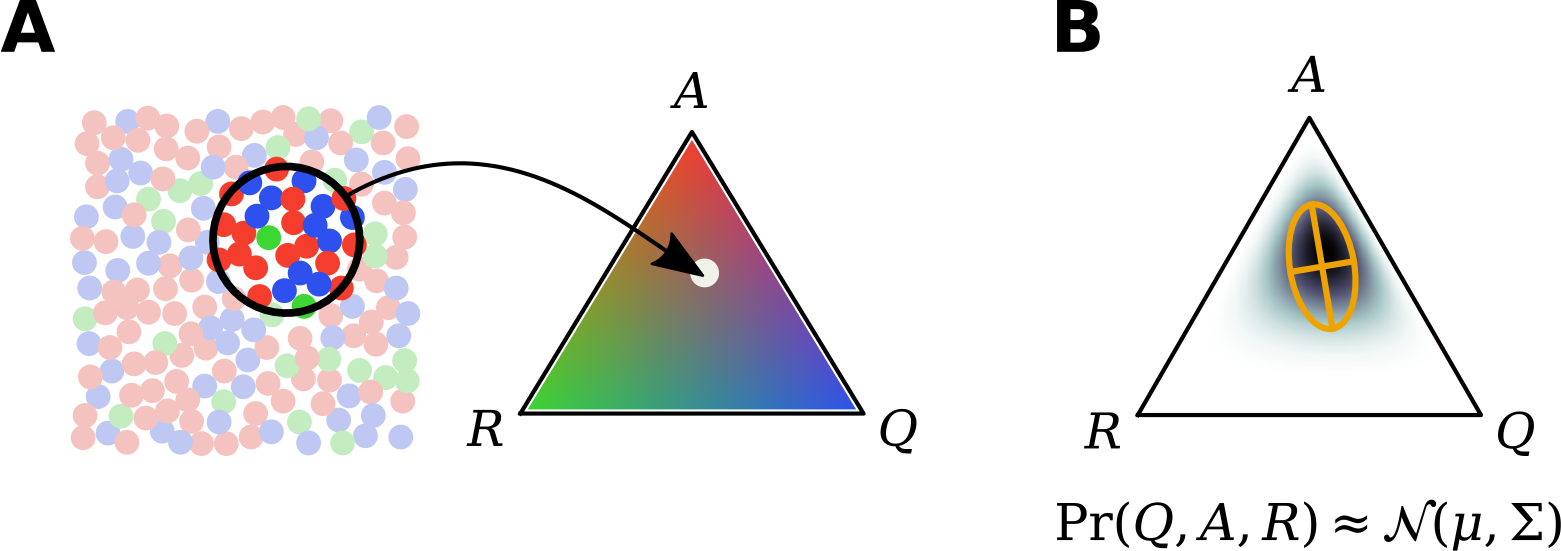
Summarizing estimated neural state as population moments. **A** The activity within a local spatial region (encircled, left) can be summarized by the fraction of cells (represented by colored dots) in the quiescent (blue), active (red), and refractory (green) states (*Q*, *A*, *R*; right). **B** An estimate of the population state can be summarized as a probability distribution Pr(*Q*, *A*, *R*) over the possible proportions of neurons in each state. A Gaussian moment-closure approximates this distribution as Gaussian, with given mean and covariance (orange crosshairs).

Applying this procedure to our system leads to the following evolution equations of the first moments (mean concentrations):

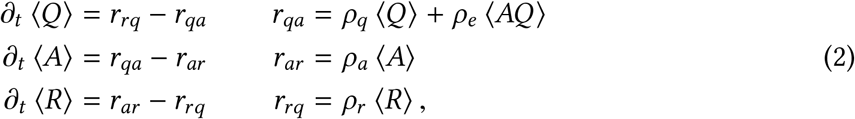

where the rate variables *r*_(·)(·)_ describe the rates of the different transitions in Eq. (1), and 〈·〉 denotes expected-value with respect to the distribution over population states. Intuitively, Eq. (2) says that the mean number of neurons in each state evolves according to the difference between the rate that neurons enter, and the rate that neurons leave, said state. For spontaneous (Poisson) state transitions, these rates are linear and depend only on the average number of neurons in the starting state. The transition from *Q* to *A*, however, has both a spontaneous and excito-excitatory component. The latter depends on the expected product of active and quiescent cells 〈*AQ*〉, which is a second moment and can be expressed in terms of the covariance: 〈*AQ*〉 = 〈*A*〉 〈*Q*〉 +Σ_*AQ*_. We obtain similar equations for the covariance of the system (Eq. 6; Methods: *Moment-closure for a single population)*. These can be solved jointly with Eq. (2) forward in time to give an approximation of the system’s dynamics.

### 2.4 Generalization to spatial (neural field) system

So far we have considered a single local population. We next extend our model to a two-dimensional spatial system. In this case the mean concentrations become density or mean fields (‘neural fields’) that depend on spatial coordinates **x** = (*x*_1_, *x*_2_, e.g. 〈*Q*〉 becomes 〈**Q**(**x**)〉. Similarly, the covariances become two-point correlation functions. For example, Σ_*QA*_(**x**, **x′**) denotes the covariance between the number of neurons in the quiescent state at location **x** and the number of neurons in the active state at location **x′**, (see Methods: *Extension to spatial system* for details).

By replacing the mean concentrations and covariances accordingly in Eqs. (2) and (6), we obtain spatial evolution equations for these space-dependent quantities. The terms arising from the linear transitions in Eq. (1) (i.e. *r*_*rq*_, *r*_*aq*_ and the first term in *r*_*qa*_ in Eq. 2) do not introduce any spatial coupling and hence do not need to be modified (note also that neurons do not diffuse or move otherwise, which is why we do not obtain a dynamic term in the resulting equations). The nonlinear excitatory interaction *Q*+*A*→*A*+*A* in Eq. (1), however, introduces a coupling which we need to specify further in a spatial setting. We assume that each quiescent neuron experiences an excitatory drive from nearby active neurons, and that the interaction strength can be described as a function of distance ║Δ**x**║ by a Gaussian interaction kernel:

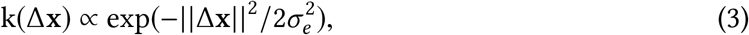

where *σ*_*e*_ the standard deviation determining the length scale of the interaction: for distances larger than *σ*_*e*_, the interaction strength decreases exponentially. This kernel introduces a spatial coupling between the neurons, which could be mediated by synaptic interactions, diffusing neurotransmitters, gap junction coupling, or combinations thereof. With this coupling, the transition rate (compare to Eq. (2)) from the quiescent to active state at position x becomes the following integral:

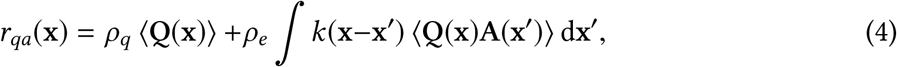

where the integral runs over the whole volume of the system. (see Methods: *Extension to spatial system* for details)

We thus obtain a ‘second-order’ neural field in terms of the mean fields and two-point correlation functions. We simulated the spatially-extended system by sampling. Figure 3 shows that it is indeed capable of producing multi-scale wave-like phenomena similar to the waves observed in the retina (c.f. Figure 5; see Methods: *Langevin equations and sampling* for details).

**Figure 3:**
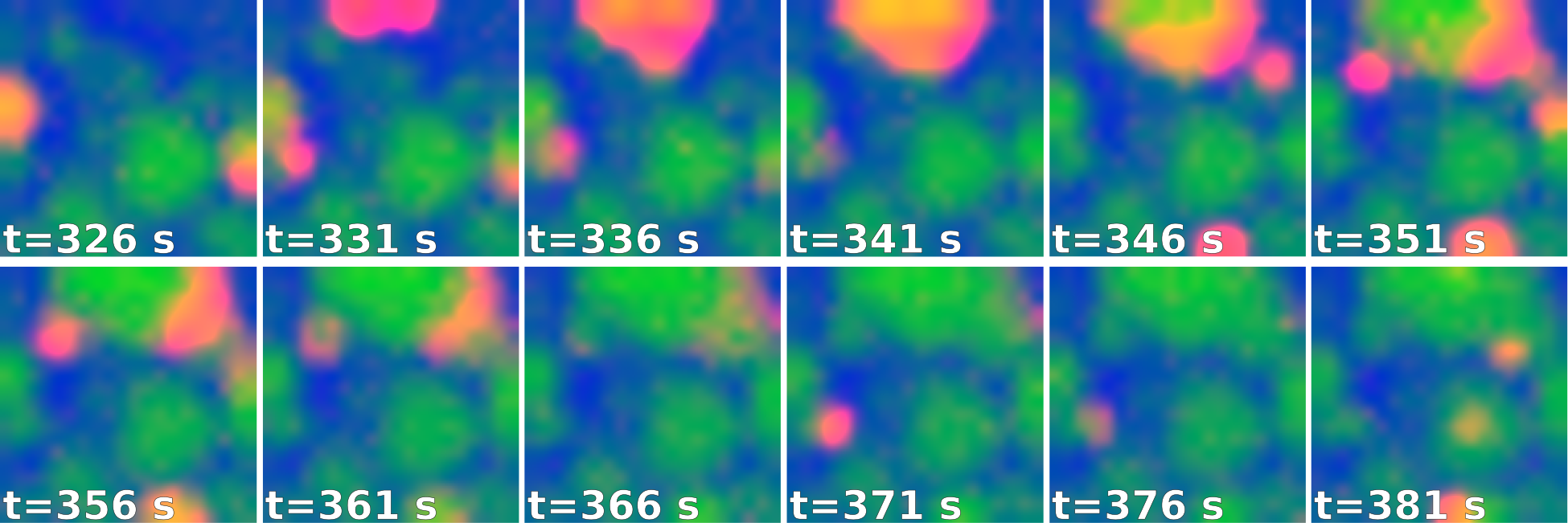
Spatial 3-state neural-field model exhibits self-organized multi-scale wave phenomena. Simulated example states at selected time-points on a [0, 1]^2^ unit interval using a 20×20 grid with effective population density of *ρ*=50 cells per unit area, and rate parameters *σ*=7.5e-2, *ρ*_*a*_=4e-1, *ρ*_*r*_=3.2e-3, *ρ*_*e*_=2.8e-2, and *ρ*_*q*_=2.5e-1 (Methods: *Langevin equations and sampling*). As, for instance, in neonatal retinal waves, spontaneous excitation of quiescent cells (blue) lead to propagating waves of activity (red), which establish localized patches in which cells are refractory (green) to subsequent wave propagation. Over time, this leads to diverse patterns of waves at a range of spatial scales.

### 2.5 Neural field models as latent-variable state-space models

The equations for the mean fields and correlations can be integrated forward in time and used as a state-space model to explain population spiking activity (Figure 4; Methods: *Bayesian flltering*).

**Figure 4:**
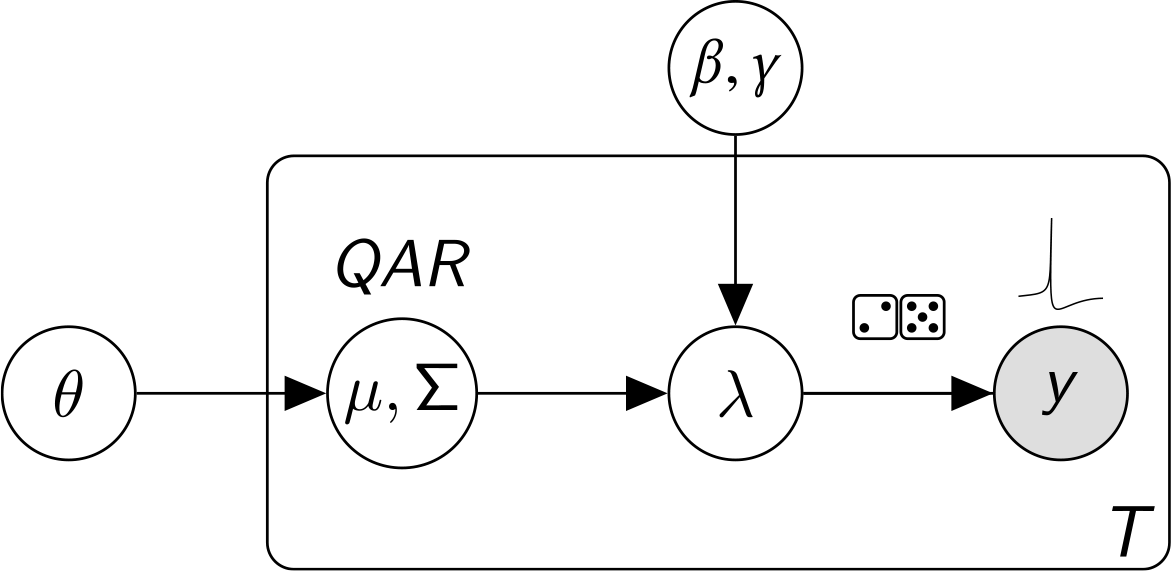
Hidden Markov model for latent neural fields. For all time-points *T*, state transition parameters *θ* = (*ρ*_*q*_, *ρ*_*a*_, *ρ*_*r*_, *ρ*_*e*_, *σ*) dictate the evolution of a multivariate Gaussian model *μ*, Σ of latent fields *Q*, *A*, *R*. The observation model (*β*, *γ*) is a linear map with adjustable gain and threshold, and reflects how field *A* couples to firing intensity *λ*. Point-process observations (spikes) *y* are Poisson with intensity *λ*.

In extracellular recordings, we do not directly observe the intensity functions 〈**Q**(**x**)〉, 〈**A**(**x**)〉, and 〈**R**(**x**)〉. Instead, we observe the spikes that active neurons emit, or in the case of developmental etinal waves recorded via a HDMEA setup, we observe the spikes of retinal ganglion cells which are driven by latent wave activity. The spiking intensity should hence depend on the density **A**(**x**) of active neurons. Here, we assume that neural firing is a Poisson conditioned on the number of active neurons, which allows us to write the likelihood of point (i.e. spike) observations in terms of **A**(**x**) (Truccolo et al. 2005, 2010; Truccolo 2016; see Methods: *Point-process measurement likelihood* for details).

The combination of this Poisson observation model with the state-space model derived in previous sections describes how hidden neural field states evolve in time and how these states drive neuronal spiking. Given spatiotemporal spiking data, the latent neural field states and correlations can then be inferred using a sequential Bayesian filtering algorithm. The latter uses the developed field model to predict the evolution of latent states, and updates this estimate at each time point based on the observed neuronal spiking (Methods: *Bayesian flltering*). This provides estimates of the unobserved physiological states of the neurons. To verify this approach, we simulated observations from the neural field equations (Figure 3; Methods: *Langevin equations and sampling*), and inferred the latent neural states and confidence intervals via Bayesian filtering using known parameters (Figure 5). We used parameters corresponding to a relatively low spike rate, indicating that state inference can recover latent states in the presence of limited measurement information.

**Figure 5:**
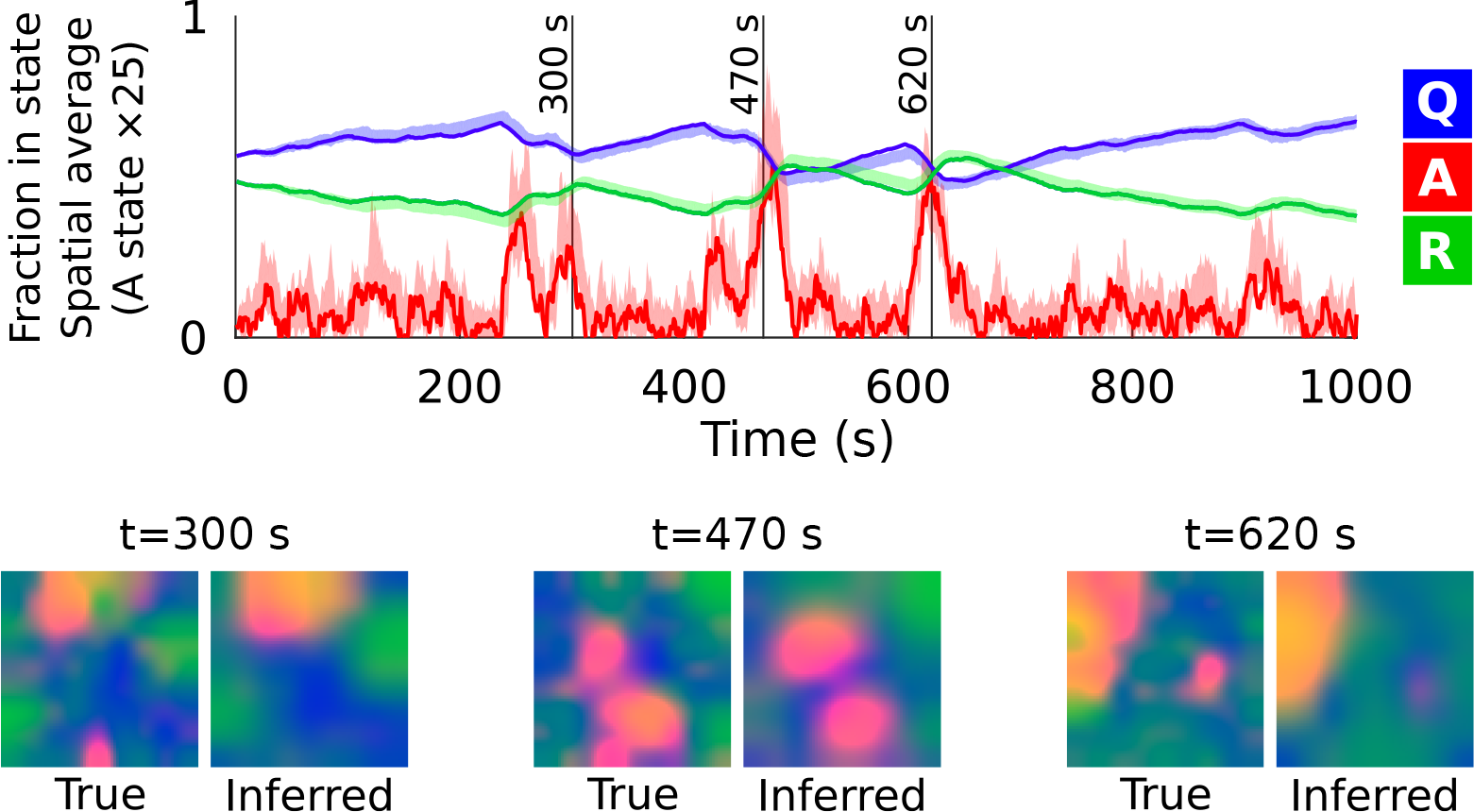
State inference via filtering: ground-truth simulation. Filtering recovers latent states in ground-truth simulated data. Spatially averaged state occupancy (**Q**, **A**, and **R**) (y-axis) is plotted over time (x-axis). Solid lines represent true values sampled from the model, and shaded regions represent the 95% confidence interval estimated by filtering. The active (**A**) state density has been scaled up by a factor of 25 for visualization. Colored plots (below) show the qualitative spatial organization of quiescent (blue), active (red), and refractory (green) neurons is recovered by filtering during example wave events. Model parameters are the same as Figure 3, with the exception of the spatial resolution, which has been reduced to a 9×9 grid. Conditionally-Poisson spikes were sampled with bias *β*=0 and gain *γ*=15 spikes/second per simulation area.

## 3 Application to retinal wave datasets

Having developed an interpretation of neural field equations as a latent-variable state-space model, we next applied this model to the analysis of spatiotemporal spiking data from spontaneous traveling wave activity occurring in the neonatal vertebrate retina (e.g. Figure 7; Sernagor et al. 2003; Hennig et al. 2009a; Blankenship et al. 2009; Meister et al. 1991; Zhou and Zhao 2000; Feller et al. 1996; Maccione et al. 2014).

**Figure 6:**
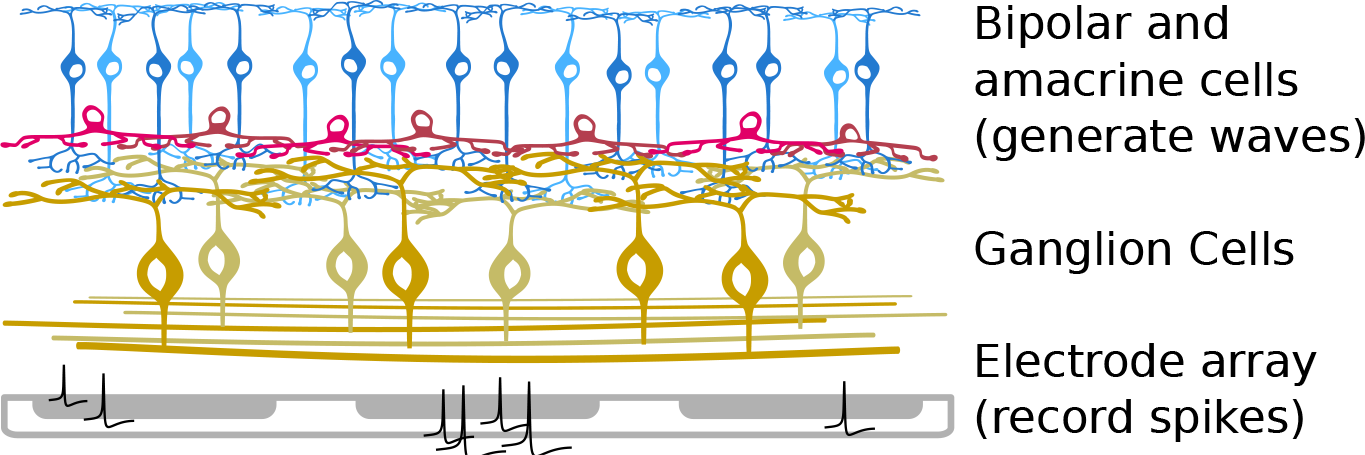
Illustration of inner retina and recording setup. Spontaneous retinal waves are generated in the inner retina via laterally interacting bipolar (blue) and amacrine (red) cells, depending on the developmental age. These waves activate Retinal Ganglion Cells (RGCs; yellow), the output cells of the retina. RGC electrical activity is recorded from the neonatal mouse retina via a 64×64 4096-electrode array with 42 μm spacing.

**Figure 7:**
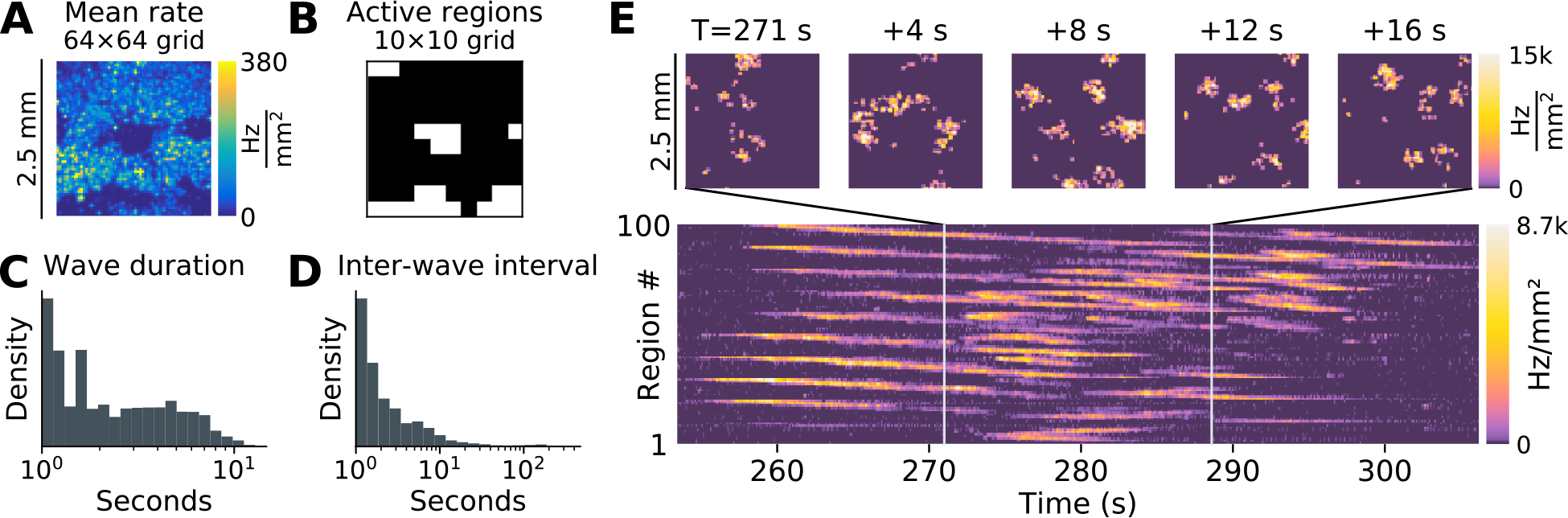
Developmental retinal waves. Example neonatal mouse retinal waves recorded on a 4096-electrode array on postnatal day 6. Recorded spikes were binned at 100 ms resolution, and assigned to 10×10 spatial regions for analysis. **A** Average firing rate from RGCs recorded across the retina (the central region devoid of recorded spikes is the optic disc). **B** Spatial regions with spiking activity detected for further analysis **C** Distribution of wave durations. To segment waves, spiking activity on each channel was segmented into “up” states (during wave activity) and “down” states (quiescent) using a two-state hidden Markov model with Poisson observations. **D** Average inter-wave interval. **E** Example wave event, traveling across multiple spatial regions and lasting for a duration of 16-20 seconds.

### 3.1 State inference in developmental retinal waves

During retinal development, the cell types that participate in wave generation change (Maccione et al., 2014; Sernagor et al., 2003; Zhou and Zhao, 2000), but the three-state model globally describes dynamics in the inner retina at all developmental stages (Figure 6). The Active (*A*) state describes a sustained bursting state, such as the depolarization characteristic of starburst amacrine cells (Figure 6) during acetylcholine-mediated early-stage (Stage 2) waves between P0 and P9 (Feller et al., 1996; Zhou and Zhao, 2000), and late-stage (Stage 3) glutamate-dependent waves (Bansal et al., 2000; Zhou and Zhao, 2000). For example, Figure 7 illustrates spontaneous retinal wave activity recorded from a postnatal day 6 mouse pup (Stage 2). In addition, at least for cholinergic waves, the slow refractory state *R* is essential for restricting wave propagation into previously active areas (Zheng et al., 2006). We note that the multi-scale wave activity exhibited in the three-state neural field model (e.g. Figure 3) recapitulates the phenomenology of retinal wave activity explored in the discrete three-state model of Hennig et al. (2009b).

Using RGC spikes recorded with a 4,096 channel HDMEA (Figure 6), we demonstrate the practicality of latent-state inference using heuristically initialized rate parameters and illustrate an example of inference for a retinal wave dataset from postnatal day 11 (Stage 3; Figure 8). For retinal wave inference, we normalize the model by population-size (Methods: *System-size scaling*) so that the gain and bias may be set independently in normalized units, rather than depending on the local neuronal population size. Model parameters were initialized heuristically based on observed timescales at *ρ*_*e*_=*ρ*_*ar*_=15, *ρ*_*r*_=0.15, and *σ* =0.15. As in Lansdell et al. (2014), lateral interactions in our model reflect an effective coupling that combines both excitatory synaptic interactions and the putative effect of diffusing excitatory neurotransmitters, which has been shown to promote late-stage glutamatergic wave propagation (Blankenship et al., 2009). The moment-closure system does not accurately approximate the rare and abruptly-discontinuous nature of wave initiation. We therefore model spontaneous wave-initiation events as an extrinsic noise source, and set the spontaneous excitation rate *ρ*_*q*_ to zero in the neural field model that defines our latent state-space. The Poisson noise was re-scaled to reflect an effective population size of 16 neurons/mm^2^, significantly smaller than the true population density (Jeon et al., 1998). However, due to the recurrent architecture and correlated neuronal firing, the effective population size is expected to be smaller than the true population size. Equivalently, this amounts to assuming supra-Poisson scaling of fluctuations for the neural population responsible for retinal waves.

**Figure 8:**
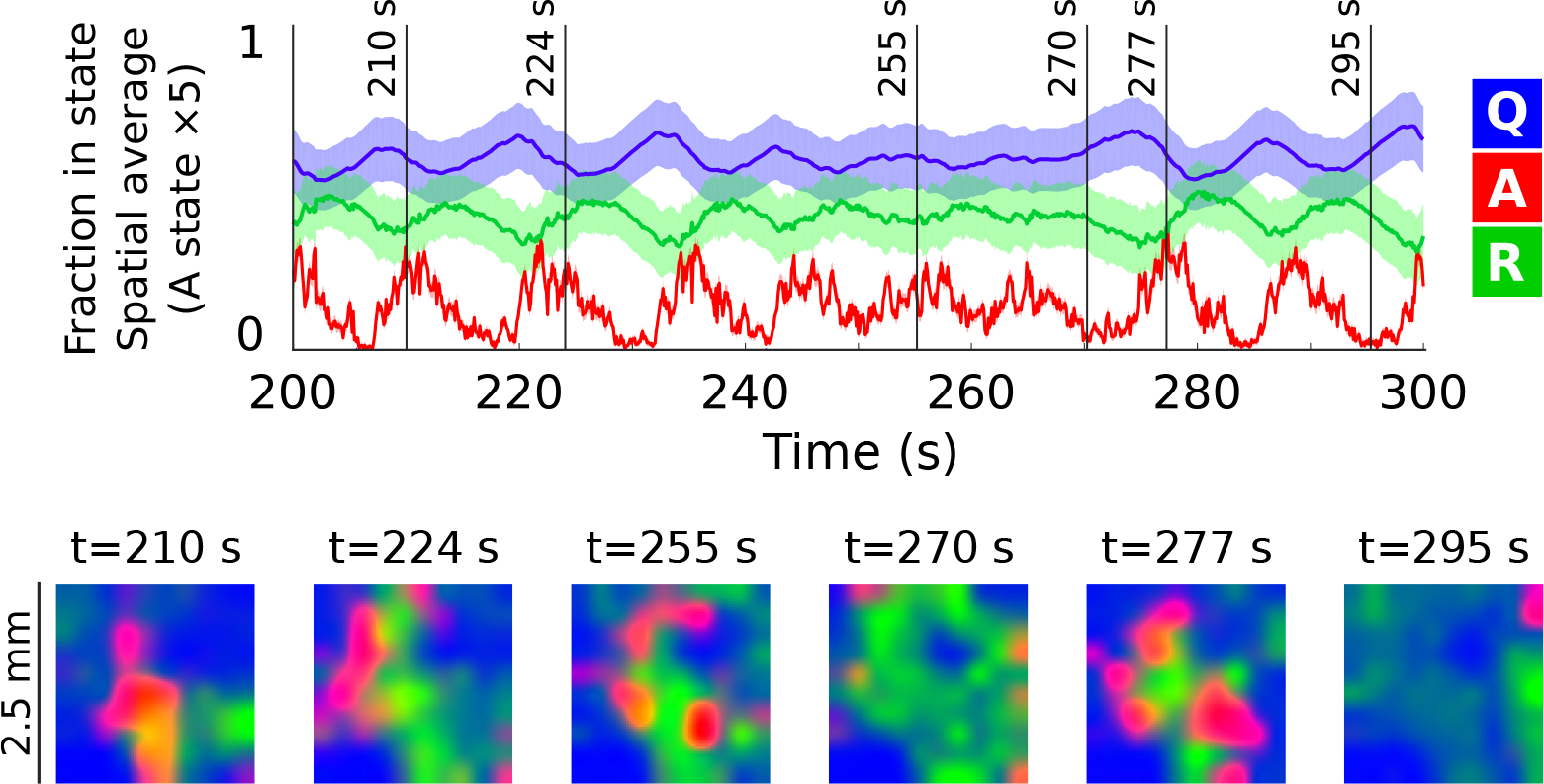
State inference via filtering: retinal datasets. *Filtering of spontaneous retinal waves (postnatal day 11).* Solid lines indicate inferred means, shaded regions the 95% confidence bound. The magnitude of the *A* state and the counts have been scaled up by a factor of 5 for visualization. Grey vertical lines indicate example time slices, which are shown in the colored plots below. Colored plots are the same as in Figure 5, with red, green, and blue reflecting (normalized) densities of active, refractory, and quiescent cells, respectively.

Bayesian filtering recovers the expected features of the retinal waves (Figure 8): the excito-excitatory transition *Q*+*A*→*A*+*A* and the onset of refractoriness *A*→*R* are rapid compared to the slow refractory dynamics, and therefore the *A* state is briefly occupied and mediates an effective *Q*→*R* transition during wave events. The second-order structure provided by the covariance is essential, as it allows us to model posterior variance (shaded regions in Figure 8), while also capturing strong anti-correlations due to the conservation of reacting agents, and the effect of correlated fluctuations on the evolution of the means. Furthermore, spatial correlations allow localized RGC spiking events to be interpreted as evidence of regional (spatially-extended) latent neuronal activity.

### 3.2 Open challenges in model identification

So far, we have demonstrated good recovery of states when the true rate parameters are known (Figure 5), and shown that plausible latent-states can be inferred from neural point-process datasets using heuristically initialized parameters (Figure 8). A natural question then is whether one can use the Bayesian state-space framework to estimate a posterior likelihood on the rate parameter values, and infer model parameters directly from data. At present, model inference remains very challenging for four reasons: under-constrained parameters, computational time complexity, numerical errors from successive approximations, and non-convexity in the joint posterior. It is worth reviewing these open challenges as they relate to important open problems in machine learning and data assimilation.

*First*, the effective population size, the typical fraction of units in quiescent vs. refractory states, and the gain parameter mapping latent activations to spiking, are all essential to setting appropriate rates, and are not accessible from observation of RGC spiking alone. Without direct measurement or appropriate physiological priors on parameter values, recovering a physiologically realistic model is infeasible. In effect, this means that a large number of equivalent systems can explain the observed RGC spiking activity, a phenomenon that has been termed “sloppiness” in biological systems (Transtrum et al., 2015; Panas et al., 2015). Indeed, Hennig et al. (2011) show that developmental waves are robust to pharmacological perturbations, suggesting that the retina itself can use different configurations to achieve similar wave patterns. *Second*, although state inference is computationally feasible, parameter inference requires many thousands of state-inference evaluations. A Matlab implementation of state-inference running on a 2.9 GHz 8-core Xeon CPU can process ∼85 samples/s for a 3-state system on a 10×10 spatial basis. For a thirty-minute recording of retinal wave activity, state inference is feasible, but repeated state inference for parameter inference is impractical. *Third*, model likelihood must be computed recursively, and is subject to well-known loss of numerical accuracy due to back-propagation through time (Pascanu et al., 2013; Bengio et al., 1994; Hochreiter et al., 2001). In other words, small errors in the past can have large effects in the future owing to the nonlinear and excitable nature of the system. This makes it difficult to dissociate numerical errors due to the recursive calculations from true estimates of the error related to model parameters. For early-stage retinal waves, the large separation of timescales between the refractory effects, and the fast-timescale excitation, also makes it difficult to estimate gradients for the slow-timescale parameters. Furthermore, the inferred likelihood is approximated as the product of a large number of high-dimensional Laplace approximations (or similar Gaussian approximations, e.g. variational), which makes the inferred model likelihood itself approximate. *Fourth* and finally, the overall likelihood surface is not, in general, convex, and may contain multiple local optima. In addition, regions of parameters space can exhibit vanishing gradient for one or model parameters. This can occur when the value of one parameter makes others irrelevant. For example, if the excito-excitatory interaction *ρ*_*e*_ is set to a low value, the interaction radius *σ*_*e*_ for excitation becomes irrelevant since the overall excitation is negligible.

Overall, parameter inference via Bayesian filtering presents a formidable technical challenge that hinges upon several open problems for efficient model inference in high-dimensional spatiotemporal point process models undergoing latent, nonlinear dynamics. At present, it would seem that traditional parameter identification methods, based on mathematical expertise and matching observable physical quantities (e.g. wavefront speed, c.f. Lansdell et al. 2014), remain the best-available approach to model estimation. Nevertheless, the state-space formulation of neural field models enables Bayesian state inference from candidate neural field models, and opens the possibility of likelihood-based parameter inference in the future.

## 4 Discussion

In this work, we showed that classical neural-field models, which capture the activity of large, interacting neural populations, can be interpreted as state-space models, where we can explicitly model microscopic, intrinsic dynamics of the neurons. This is achieved by interpreting a second-order neural field model as defining equations on the first two moments of a latent-variable process, which is coupled to spiking observations. In the state-space model interpretation, latent neural field states can be recovered from Bayesian filtering. This allows inferring the internal states of neuronal populations in large networks based solely on recorded spiking activity, information that can experimentally only be obtained with whole cell recordings. We demonstrated successful state inference for simulated data, where the correct model and parameters were known. Next, we applied the model to large-scale recordings of developmental retinal waves. Here the correct latent state model is unknown, but a relatively simple three-state model with slow refractoriness is well motivated by experimental observations (Zheng et al., 2006). Consistent with previous work (Feller et al., 1997; Zheng et al., 2006; Godfrey and Swindale, 2007; Hennig et al., 2009a), the state inference revealed that activity-dependent refractoriness restricts the spatial spreading of waves. In contrast to phenomenological latent state-space models, the latent states here are motivated by an (albeit simplified) description of single-neuron dynamics, and the state-space equations arise directly from considering the evolution of collective activity as a stochastic process.

In the example explored here, we use Gaussian moment-closure to arrive at a second-order approximation of the distribution of latent states and their evolution. In principle, other distributional assumptions may also be used to close the moment expansion. Other mathematical approaches that yield second-order models could also be employed, for example the linear noise approximation (Van Kampen, 1992), or defining a second cumulant in terms of the departure of the model from Poisson statistics (Buice et al., 2010). The approach applied here to a three-state system can generally be applied to systems composed of linear and quadratic state transitions. Importantly, systems with only linear and pairwise (quadratic) interactions can be viewed as a locally-quadratic approximation of a more general smooth nonlinear system (Ale et al., 2013), and Gaussian moment closure therefore provides a general approach to deriving approximate state-space models in neural population dynamics.

The state-space interpretation of neural field models opens up future work to leverage the algorithmic tools of SSM estimation for data assimilation with spiking point-process datasets. However, challenges remain regarding the retinal waves explored here, and future work is needed to address these challenges. Model likelihood estimation is especially challenging. Despite this, the connection between neural-field models and state-space models derived here will allow neural field modeling to incorporate future advances in estimating recursive, nonlinear, spatiotemporal models. We also emphasize that some of the numerical challenges inherent to high-dimensional spatially extended neural field models do not apply to simpler, low-dimensional neural mass models, and the moment-closure framework may therefore provide a practical avenue to parameter inference in such models.

In summary, this report connects neural field models, which are grounded in models of stochastic population dynamics, to latent state-space models for population spiking activity. This connection opens up new approaches to fitting neural field models to spiking data. We expect that this interpretation is a step toward the design of coarse-grained models of neural activity that have physically interpretable parameters, have physically measurable states, and retain an explicit connection between microscopic activity and emergent collective dynamics. Such models will be essential for building models of collective dynamics that can predict the effects of manipulations on single-cells on emergent population activity.

## Acknowledgements

Funding provided by EPSRC EP/L027208/1 *Large scale spatiotemporal point processes: novel machine learning methodologies and application to neural multi-electrode arrays*. We thank Gerrit Hilgen for important discussions in establishing biologically-plausible parameter regimes for the three-state model. We thank Evelyne Sernagor for the retinal wave datasets, as well as ongoing advice and invaluable feedback on the manuscript.

## 5 Methods

### 5.1 Data acquisition and preparation

Example retinal wave datasets are taken from Maccione et al. (2014). Spikes were binned at 100 ms resolution for analysis, and regions without spiking observations were excluded. Spiking activity in each region was segmented into wave-like and quiescent states using a two-state hidden Markov model with a Poisson observations. To address heterogeneity in the Retinal Ganglion Cell (RGC) outputs, the observation model was adapted to each spatial region based on firing rates. Background activity was used to establish per-region biases, defined as the mean activity in a region during quiescent periods. The scaling between latent states and firing rate (gain) was adjusted locally based on the mean firing rate during wave events. The overall (global) gain for the observation model was then adjusted so that 99% of wave events in a given region corresponded to a fraction of cells in the active (*A*) state fraction no greater than one.

### 5.2 Moment-closure for a single population

To develop a state-space formalism for inference and data assimilation in neural field models, we begin with a master equation approach. This approach has been used before to analyze various stochastic neural population models, often as a starting point to derive ordinary differential equations for the moments of the distribution of population states, as we do here (Ohira and Cowan, 1993; Buice and Cowan, 2007; Bressloff, 2009; El Boustani and Destexhe, 2009; Buice et al., 2010; Touboul and Ermentrout, 2011). In our case, we examine a three-state system of the kind proposed in Buice and Cowan (2007, 2009), and use a Gaussian moment-closure approach similar to Bressloff (2009). The master equation describes how the joint probability distribution of neural population states (in our example the active, quiescent and refractory states) evolves in time. However, modelling this full distribution is computationally prohibitive for a spatially-extended system, since the number of possible states scales exponentially with the number of neural populations. Instead, we approximate the time evolution of the moments of this distribution. In principle, an infinite number of moments are needed to describe the full population activity. To limit this complexity, we consider only the first two moments (mean and covariance), and use a moment-closure approach to close the series expansion of network interactions in terms of higher moments (Schnoerr et al. 2017; Gomez-Uribe and Verghese 2007; Goodman 1953; Whittle 1957; for applications to neuroscience see Ly and Tranchina 2007; Bressloff 2009; El Boustani and Destexhe 2009; Buice et al. 2010; Touboul and Ermentrout 2011; Rule and Sanguinetti 2018). Using this strategy, we obtain a second-order neural field model that describes how the mean and covariance of population spiking evolve in time, and recapitulates spatiotemporal phenomena when sampled.

We may describe the number of neurons in each state in terms of a probability distribution Pr(*Q*, *A*, *R*) (Figure 2A), where we slightly abuse notation and use *Q*, *A*, and *R* both as symbols for the neuron states and as variables counting the neurons in the corresponding states, i.e. non-negative integers. The time evolution of this probability distribution captures stochastic population dynamics, and is represented by a master equation that describes the change in density for a given state {*Q*, *A*, *R*} when neurons change states. Accordingly, the master equation describes the change in probability of a given state {*Q*, *A*, *R*} in terms of the probability of entering, minus the probability of leaving the state:

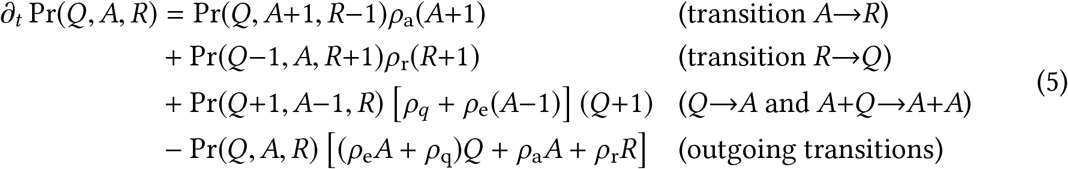

Even in this simplified non-spatial scenario, no analytic solutions are known for the master equation. However, from Eq. (5) one can derive equations for the mean and covariance of the process. The approach, generally, is to consider expectations of individual states, e.g. 〈*Q*〉 (first moments, i.e. means), or 〈*QA*〉 (second moments), taken with respect to the probability distribution Pr(*Q*, *A*, *R*) described by the master equation (5). Differentiating these moments in time, and substituting in the time-evolution of the probability density as given by the master equation, yields expressions for the time-evolution of the moments. However, in general these expressions will depend on higher moments and are therefore not closed.

For our system, the nonlinear excitatory interaction *Q*+*A*→*A*+*A* couples the evolution of the means to the covariance Σ_*AQ*_, and the evolution of the covariance is coupled to the third moment, and so on. The moment equations are therefore not closed, and require an infinite number of moments to describe the evolution of the mean and covariance. To address this computational complexity, we approximate Pr(*Q*, *A*, *R*) with a multivariate normal distribution at each time-point (Figure 2B), thereby replacing counts of neurons with continuous variables. This Gaussian moment-closure approximation sets all cumulants beyond the variance to zero, yielding an expression for the third moment in terms of the mean and covariance, leading to closed ordinary differential equations for the means and covariances (Goodman, 1953; Whittle, 1957; Gomez-Uribe and Verghese, 2007; Schnoerr et al., 2017).

For out model with transitions given in Eq. (1) this leads to the system of ODEs for the mean values given in Eq. (2) in the main text. For the evolution of the covariance we obtain

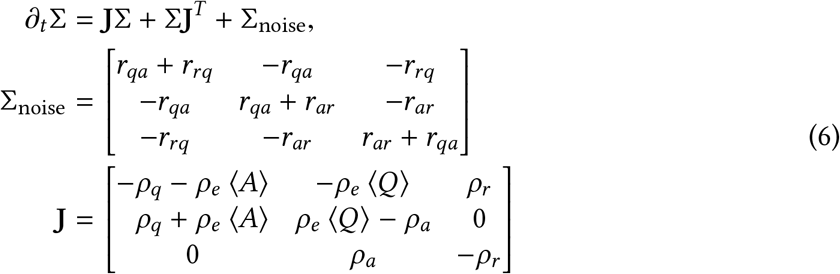

where **J** is the Jacobian of the equations for the deterministic means in Eq. (2), and the Σ_noise_ fluctuations are Poisson and therefore proportional to the mean reactions rates (Eq. (2)). Intuitively, the Jacobian terms **J** describe how the covariance of the state distribution ‘stretches’ or ‘shrinks’ along with the deterministic evolution of the means, and the additional Σ_noise_ reflects added uncertainty due to the fact that state transitions are stochastic. Each state experiences Poisson fluctuations with variance equal to the mean transition rates, due to the sum of transitions into and away from the state. Because the number of neurons is conserved, a positive fluctuation into one state implies a negative fluctuation away from another, yielding off-diagonal anticorrelations in the noise.

Together, equations (2) and (6) provide approximate equations for the evolution of the first two moments of the master equation (Eq. 5), expressed in terms of ordinary differential equations governing the mean and covariance of a multivariate Gaussian distribution. Here, we have illustrated equations for a 3-state system, but the approach is general and can be applied to any system with spontaneous and pairwise state transitions.

### 5.3 Extension to spatial system

To extend the moment equations (2) and (6) to a neural field system, we consider a population of neurons at each spatial location. In this spatially-extended case, we denote the intensity fields as **Q**, **A**, and **R**, which are now vectors with spatial indices (or, in the spatially-continuous case: scalar functions of coordinates **x**). In the spatially-extended system, active (**A**) neurons can excite nearby quiescent (**Q**) neurons. We model the excitatory influence of active cells as a weighted sum over active neurons in a local neighborhood, defined by a coupling kernel *K*(Δ**x**) that depends on distance (Eq. 4). To simplify the derivations that follow, denote the convolution integral in equation (4) as a linear operator **K** such that

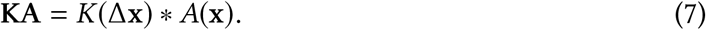

In this notation, one can think of **K** as a matrix that defines excitatory coupling between nearby spatial regions. Using the notation of Eq. (7), the rate that active cells excite quiescent ones is given by the product

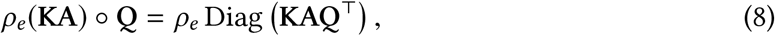

where ο denotes element-wise (in the spatially-continuous case: function) multiplication. For the time evolution of the first moment (mean intensity) of **Q** in the spatial system, one therefore considers the expectation 〈KAQ^⊤^〉, as opposed to 〈*AQ*〉 in the non-spatial system. Since **K** is a linear operator, and the extension of the Gaussian state-space model over the spatial domain **x** is a Gaussian process, the second moment of the nonlocal interactions **KA** with **Q** can be obtained in the same way as one obtains the correlation for a linear transformation of a multivariate Gaussian variable:

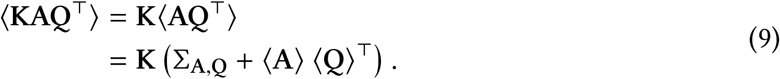

The resulting equations for the spatial means are similar to the nonspatial system (Eq. 2), with the exception that we now include spatial coupling in the rate at which quiescent cells enter the active state:

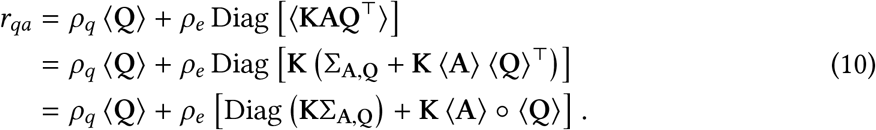

The number of neurons in the quiescent verses active states are typically anti-correlated, because a neuron entering the active state implies that one has left the quiescent state. Therefore, the expected number of interactions between quiescent and active neurons is typically smaller than what one might expect from the deterministic mean field alone. The influence of correlations Diag (**K**Σ_A,Q_) on the excitation is therefore important for stabilizing the excitatory dynamics.

To extend the equations for the second moment to the neural field case, we consider the effect of spatial couplings on the the Jacobian (Eq. 6). The spontaneous first-order reactions remain local, and so the linear contributions are similar to the non-spatial case. However, nonlocal interaction terms emerge in the nonlinear contribution to the Jacobian:

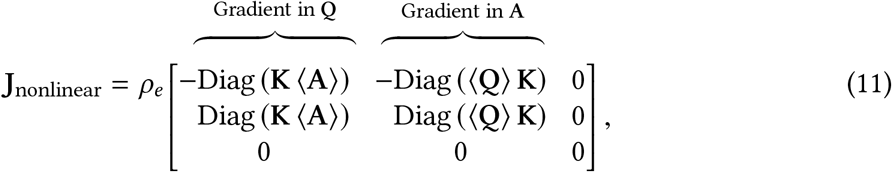

where here the “Diag” operation refers to constructing a diagonal matrix from a vector. Intuitively, the first column of Eq. (11) reflects the fact that the the availability of quiescent cells modulates the excitatory effect of active cells, and the second column reflects the fact that the density active of neurons in nearby spatial volumes contribute to the rate at which quiescent cells become active.

### 5.4 Basis projection

The continuous neural field equations are simulated by projection onto a finite spatial basis *B*. Each basis element is an integral over a spatial volume. Means for each basis element are defined as an integral over this volume, and correlations are defined as a double integral. For example, consider the number of quiescent neurons associated with the *i*^*th*^ basis function, *Q*_*i*_. The mean 〈*Q*_*i*_〉 and covariance 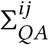 between the quiescent and active states are given by the projections:

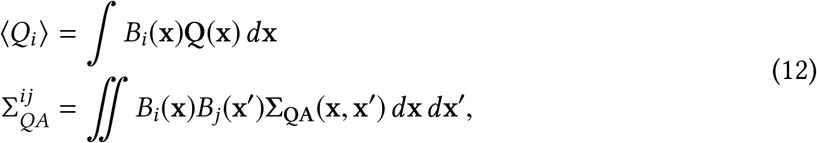

where **x** and **x′**, range over spatial coordinates as in Eq. (3) and (4). When selecting a basis *B*, assumptions must be made about the minimum spatial scale to model. A natural choice is the radius of lateral (i.e. spatially nonlocal) interactions in the model *σ*_*e*_ (Eq. 3), since structure below this scale is attenuated by the averaging over many nearby neurons in the dendritic inputs.

### 5.5 Langevin equations and sampling

For ground-truth simulations, we sample from a hybrid stochastic model derived from a Langevin approximation to the three-state neural field equation. In the Langevin approximation, the deterministic evolution of the state is given by the mean-field equations (Eq. (2) for a local system, Eq. (10) for the neural field system), and the stochastic noise arising from Poisson state transitions is approximated as Gaussian as given by second-order terms (i.e. Σ_noise_ in Eq. (6); see also Riedler and Buckwar, 2013; Schnoerr et al., 2017). Spontaneous wave initiation events are too rare to approximate as Gaussian, and instead are sampled as Poisson (shot) noise, giving us a hybrid stochastic model:

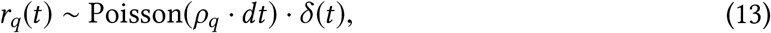

where *δ*(*t*) is a Dirac delta (impulse). To avoid uniform spontaneous excitation, the excito-excitatory reaction rate is adjusted by a small finite threshold *ϑ*, i.e. *r*_*qa*_ ← max(0, *r*_*qa*_ −*ϑ*) in Eq. (10).

For our simulations (e.g. Figure 3), we let *ϑ*=8e-3. For the non-spatial system, the hybrid stochastic differential equation equation is:

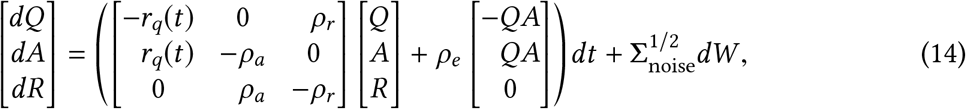

where Σ_noise_ is the fluctuation noise covariance as in Equation (6) (with *ρ*_*q*_ excluded, as it is addressed by the shot noise, Equation (13)), and *dW* is the derivative of a multidimensional standard Wiener process, i.e. a spherical (white) Gaussian noise source. The deterministic component of the Langevin equation can be compared to Equation (2) for the means of the non-spatial system in the moment-closure system (without the covariance terms).

The stochastic differential equation for the spatial system is similar, consisting to a collection of local populations coupled through the spatial interaction kernel (Eqs. 3-4), and follows the same derivation used when extending the moment-closure to the spatial case (Methods: *Extension to spatial system*, Eqs. 7-10). When applying the Euler-Maruyama method method to the spa-tiotemporal implementation, fluctuations were scaled by 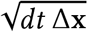, where Δ**x** is the volume of the spatial basis functions used to approximate the spatial system (See Methods: *System-size scaling* for further detail). The Euler-Maruyama algorithm samples noise from a Gaussian distribution, and can therefore create negative intensities due to discretization error. We addressed this issue by using the complex chemical Langevin equation (Schnoerr et al., 2014), which accommodates transient nonphysical negative states.

### 5.6 Point-process measurement likelihood

Similarly to generalized linear point-process models for neural spiking (Truccolo et al., 2005, 2010; Truccolo, 2016), we model spikes as a Poisson process conditioned on a latent intensity function *λ*(**x**, *t*), which characterises the probability of finding a certain number of spikes in a small spatiotemporal interval Δ*x*×Δ*t* as:

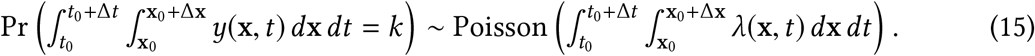

In (15), *y*(**x**, *t*) denotes the experimentally-observed spiking output, and is a sum over Dirac delta distributions corresponding to each spike with an associated time *t*_*i*_ and spatial location **x**_*i*_, i.e. *y*(**x**, *t*) = ∑_*i*∈1‥*N*_ *δ*(**x**_*i*_)*δ*(*t*_*i*_). We use a linear Poisson likelihood for which the point-process intensity function

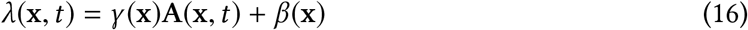

depends linearly on the number of active neurons **A**(**x**, *t*) with spatially-varying gain *γ*(**x**) and bias *β*(**x**). In other words, the observed firing intensity in a given spatiotemporal volume should be proportional to the number of active neurons, with some additional offset or bias *β* to capture background spiking unrelated to the neural-field dynamics.

### 5.7 Bayesian filtering

Having established an approach to approximate the time-evolution of the moments of a neural field system, we now discuss how Bayesian filtering allows us to incorporate observations in the estimation of the latent states. Suppose we have measurements *y*_0_, …, *y*_*N*_ of the latent state *x* at time *t*_0_, … , *t*_*N*_ given by a measurement process 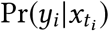, which in our case is given by the point-process likelihood (Eq. 16). Bayesian filtering allows us to recursiveely estimate the *flltering distribution* 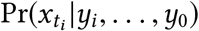 at time *t*_*i*_, i.e. the posterior state probability at time *t*_*i*_ given the current and all previous observations. The procedure works by the following iterative scheme: i) suppose we know the filtering distribution 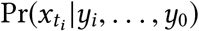 at time *t*_*i*_. Solving the dynamics forward in time up to *t*_*i*+1_ gives the predictive distribution Pr(*x*_*t*_|*y*_*i*_, … , *y*_0_) for all times *t*_*i*_<*t*≤*t*_*i*+1_. ii) at the time *t*_*i*+1_ the measurement *y*_*i*+1_ needs to be taken into account which can be done by means of the Bayesian update:

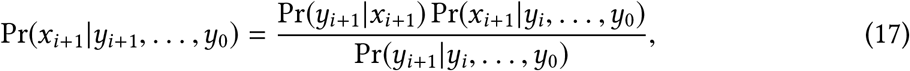

where we have used the Markov property and Pr(*y*_*i*+1_|*x*_*i*+1_, *y*_*i*_, …, *y*_0_) = Pr(*y*_*i*+1_|*x*_*i*+1_) to obtain the right hand side. Eq. (17) gives the filtering 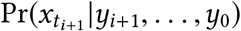 at time *t*_*i*+1_ which serves as the input of the next *i* step. Performing steps i) and ii) iteratively hence provides the filtering distribution for all times *t*_0_ ≤ *t* ≤ *t*_*n*_.

For our neural field model we must compute both steps approximately: to obtain the predictive distribution in step i) we integrate forward the differential equations for mean and covariance derived from moment-closure (Eq. 2-6 and Methods: *Extension to spatial system*). In practice, we convert the continuous-time model to discrete time. If *F*_∂*t*_ denotes the local linearization of the mean dynamics in continuous time such that ∂_*t*_ *μ*(*t*) = *F*_∂*t*_ *μ*(*t*), then the approximated discrete-time forward operator is

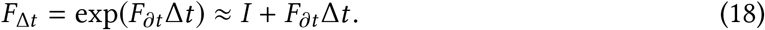

We update the covariance using this discrete-time forward operator, combined with an Euler integration step for the Poisson fluctuations. A small constant diagonal regularization term Σ_reg_ can be added, if needed, to improve stability. The resulting equations read:

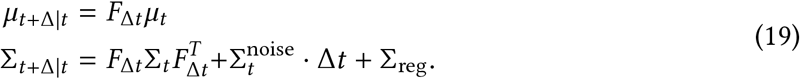

This form is similar to the update for a discrete-time Kalman filter (Kalman et al., 1960; Kalman and Bucy, 1961), the main difference being that the dynamics between observation times are taken from the nonlinear moment equations.

Consider next the measurement update of step ii) in Eq. (17). Since the Gaussian model for the latent states *x* is not conjugate with the Poisson distribution for observations *y*, we approximate the posterior Pr(*x*_*i*+1_|*y*_*i*+1_, *y*_*i*_, …, *y*_0_) using the Laplace approximation (c.f. Paninski et al. 2010; Macke et al. 2011). The Laplace-approximated measurement update is computed using a Newton-Raphson algorithm. The measurement update is constrained to avoid negative values in the latent fields by adding a *ε/x* potential (compare to the log-barrier approach; Nazarpour et al. 2012), which ensures that the objective function gradient points away from this constraint boundary, where *x* is the intensity of any of the three fields. The gradients and Hessian for the posterior measurement log-likelihood ln 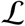 are

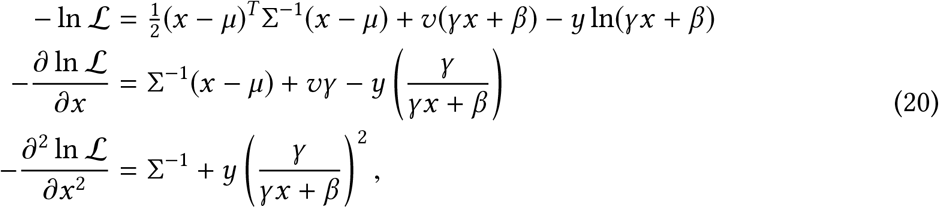

where *x* is the latent state with prior mean *μ* and covariance Σ, and couples to point-process observations *y* linearly with gain *γ* and bias *β* as in Eq. (16). The parameter *v*=Δ*x*^2^·Δ*t* is the spatiotemporal volume of the basis function or spatial region over which the counts are observed.

### 5.8 System-size scaling

For clarity, the derivations in this paper are presented for a population of neurons with a known size, such that the fields **Q**(**x**), **A**(**x**), and **R**(**x**) have units of *neurons*. In practice, the population size Ω of neurons is unknown, and it becomes expedient to work in normalized intensities, where **Q**(**x**), **A**(**x**), and **R**(**x**) represent the *fraction* of neurons in a given state between 0 and 1, and are constrained such that **Q**(**x**)+**A**(**x**)+**R**(**x**)=1. In this normalized model for population size Ω, quadratic interaction parameters (like *ρ*_*e*_) as well as the gain are multiplied by Ω, to reflect the re-scaled population. In contrast, noise variance should be *divided* by Ω to account for the fact that the coefficient of variation decreases as population size increases. Although rescaling by Ω is well-defined for finite-sized populations, the infinitesimal neural-field limit for the second-order model is not. This is because, while the mean-field equations scale with the population size 𝒪(Ω), the standard deviation of Poisson fluctuations scales with the square root of the population size 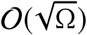. The ratio of fluctuations to the mean (coefficient of variation) therefore scales as 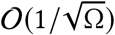, which diverges as Ω→0.

This divergence is not an issue in practice as all numerical simulations are implemented on a set of basis functions with finite nonzero volumes, and each spatial region is therefore associated with finite nonzero population size. Even in the limit where fluctuations would begin to diverge, one can treat the neural field equations as if defined over a continuous set of overlapping basis functions with nonzero volume. Conceptually, one can interpret this as setting a minimum spatial scale for the neural field equations, defined by spatial extent of each local population. If one defines the model over a set of overlapping spatial regions, then overlapping spatial areas also experience correlated fluctuations. Consider Poisson fluctuations as entering with some rate-density *σ*^2^(**x**) per unit area. The observed noise variances and covariances, projected onto basisfunctions *B*_*i*_(**x**) and *B*_*j*_(**x**), are:

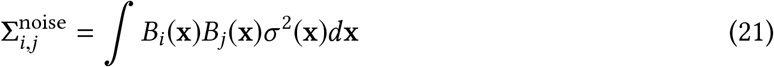

If the neuronal population density is given as *ρ*(**x**) per unit area, then the effective population size for a given basis function is:

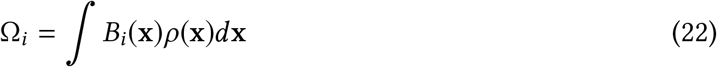

If the population density is uniform, and if basis functions have a constant volume *v*, we can write this more simply as Ω = *vρ*. In the system-size normalized model, the contributions of basis function volume cancel and the noise variance should be scaled simply as 1/*ρ*.

